# Protein Dynamics Govern the Oxyferrous State Lifetime of an Artificial Oxygen Transport Protein

**DOI:** 10.1101/2023.06.09.544418

**Authors:** Lei Zhang, Mia C. Brown, Andrew C. Mutter, Kelly N. Greenland, Jason W. Cooley, Ronald L. Koder

## Abstract

It has long been known that the alteration of protein side chains which occlude or expose the heme cofactor to water can greatly affect the stability of the oxyferrous heme state. Here we demonstrate that the rate of dynamically-driven water penetration into the core of an artificial oxygen transport protein also correlates with oxyferrous state lifetime by reducing global dynamics, without altering the structure of the active site, via the simple linking of the two monomers in a homodimeric artificial oxygen transport protein using a glycine-rich loop. The tethering of these two helices does not significantly affect the active site structure, pentacoordinate heme binding affinity, reduction potential, or gaseous ligand affinity. It does, however, significantly reduce the hydration of the protein core as demonstrated by resonance Raman spectroscopy, backbone amide hydrogen exchange, and pKa shifts in buried histidine side chains. This further destabilizes the charge-buried entatic state and nearly triples the oxyferrous state lifetime. These data are the first direct evidence that dynamically-driven water penetration is a rate-limiting step in the oxidation of these complexes. It furthermore demonstrates that structural rigidity which limits water penetration is a critical design feature in metalloenzyme construction and provides an explanation for both the failures and successes of earlier attempts to create oxygen-binding proteins.

**Significance:** This communication sheds light on one of the more controversial areas in protein folding and design: the dynamic nature of the hydrophobic core and its relationship to metalloprotein function, in particular the relationship between dynamic solvent penetration into the protein core and the stability of metalloenzyme intermediates. We demonstrate that the basic tetrameric scaffold that is the classic helical bundle model for cofactor binding and activation can be easily upgraded to a more rigid, less dynamic, single chain helical bundle by merely taking the same helical sequences and converting it to a single chain protein connected by simple, nonoptimized glycine-rich loops. Importantly, our results explain the decades-long history of failure in the design of proteins capable of stably forming an oxyferrous state – the requirement for a protein large enough to protect the heme porphyrin surface with both structural specificity and sufficient structural rigidity to restrict water penetration into the protein core. Finally, we believe this is the first use of Deep UV Resonance Raman spectroscopy to monitor dynamic water penetration in a functional protein. This method may prove useful moving forward to many research groups.

## Introduction

The hallmark of hexacoordinate hemoglobins is that they have two ligands in the oxidized state, but in the reduced state, they exist in a mixed hexa- and pentacoordination state in which one of the ligands – termed the distal ligand - has a weak affinity for the ferrous heme iron (1,2). The partial pentacoordination of the heme cofactor allows for the binding of molecular oxygen at the ferrous iron. In the case of the artificial oxygen transport protein HP7 this destabilization is the result of the coupling of histidine side chain ligation with the burial of three charged glutamate residues (red triangles, Figure 1C) on the same helix, creating a high energy ‘entatic’ conformational state (3–6). Relief of the strain of buried charges by the rotation of the glutamates into solution necessitates the detachment of the adjacent histidine side chain.

**Figure 1.**
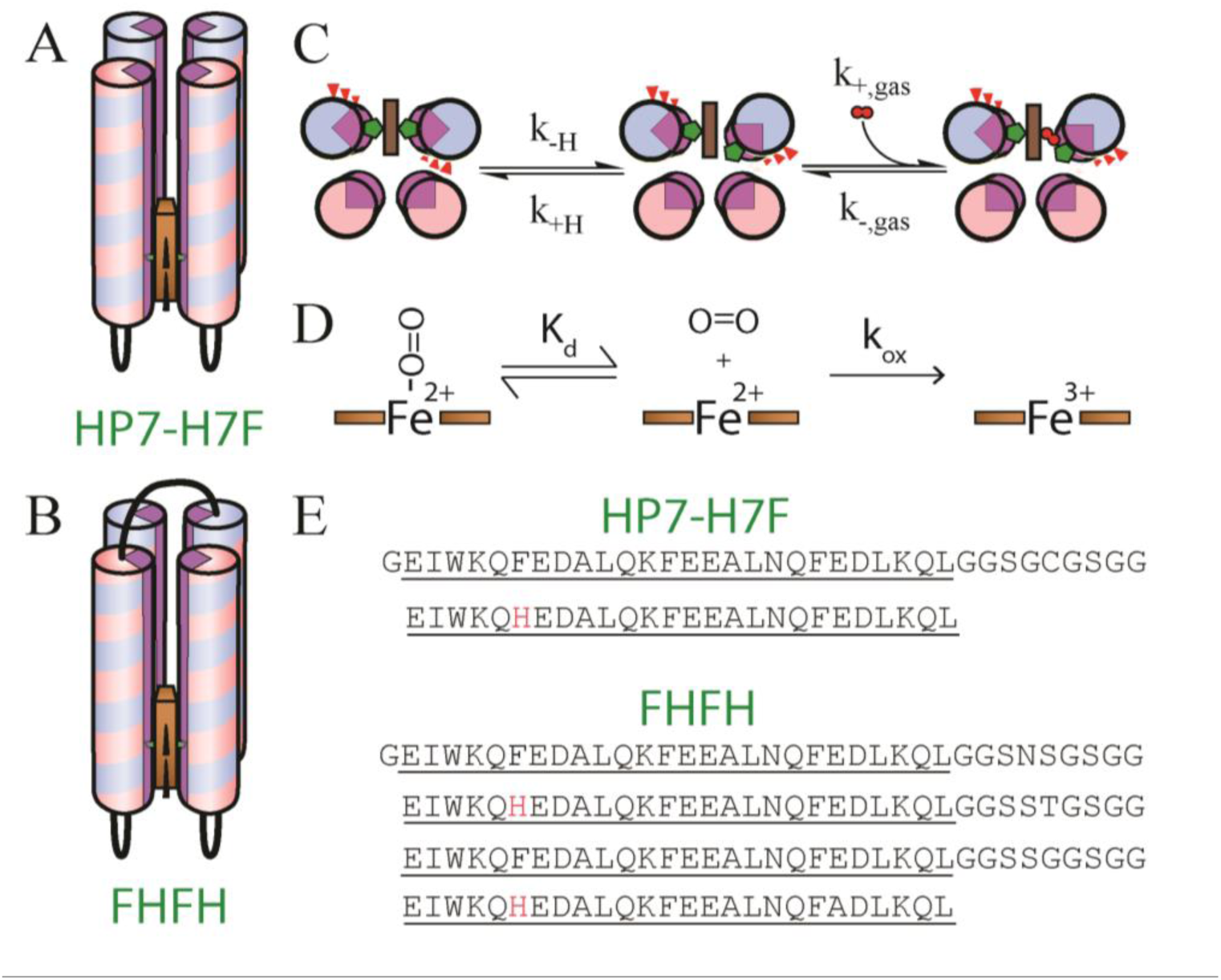
Structure, sequence and oxygen binding mechanism of two artificial oxygen transport proteins. Topology of the homodimeric protein HP7-H7F (A) and the single-chain protein FHFH (B). (C) Mechanism of oxygen binding to the ferrous heme. (D) Equilibrium and rate constants of the oxidation of the oxyferrous state. (E) Sequences of the two proteins.

Oxygen binding by heme proteins is complicated by the requirement that the heme iron be in the ferrous state for oxygen ligation. When exposed to oxygen, the ferrous heme in oxygen transport proteins can be oxidized by collisional electron exchange when not bound to molecular oxygen, but once formed, the oxyferrous complex has a reduction potential too low to be oxidized by a second molecule of oxygen (Figure 1D) (7). As we have shown (4), the rate of oxidation of the oxyferrous state is thus a function of the total protein concentration, the total oxygen concentration, the oxygen dissociation constant K_d_, and the rate constant for the reaction between oxygen and the ferrous heme complex, k_ox_:

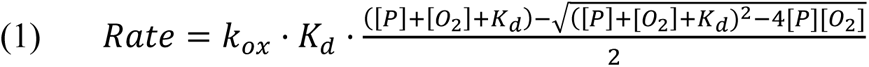

Oxidation of the oxyferrous complex in model systems is greatly accelerated by water (8–10). This is best demonstrated by the fact that imidazole-ligated hemes embedded in non-polar plastics stably form oxyferrous states with several-minute lifetimes, while the same systems dissolved in water rapidly oxidize, forming superoxide (11,12). Similarly, mutations of active site residues in bovine myoglobin which sterically increase or decrease the solvent exposure of the bound heme cofactor decrease and increase, respectively, oxyferrous state lifetimes by over two orders of magnitude, and this was shown to be of a concerted reaction consisting of the protonation of the bound oxygen by water and the dissociation of neutral superoxide (9). However, water penetration to buried heme sites in proteins driven by sidechain and backbone dynamics has not yet been demonstrated to affect lifetimes. Dynamic effects have been suggested to play a role by computational simulation of the autooxidation of myoglobin (13), and Olson and coworkers have shown that dilute preparations of human hemoglobin dissociate into dimers which oxidize more than 10-fold more quickly than the native tetramer (14). However, neither the conformational distribution nor the water penetration of the latter have been characterized.

The artificial diheme protein HP7 has proven to be a useful model system, or protein “maquette” (15), for the study of natural oxygen transport proteins. In an examination of a simplified version of the protein containing only one heme binding site, we have demonstrated that the mutational removal of all three buried glutamate side chains from the remaining heme-binding helices in HP7-H7F results in stronger distal histidine affinity, which strengthens heme cofactor binding (15,16), but weakens carbon monoxide binding by competition. However, the mutant protein does not stably bind oxygen. This loss of function was attributed to either or both of two observable effects of the mutation: the 22-fold decrease in the rate of detachment of the distal histidine ligand, which results in a longer time period for the bis-histidine bound ferrous heme to be collisionally oxidized by free molecular oxygen before the oxyferrous state can be formed, or the greatly increased degree of water penetration into the protein core, evinced by the fact that hydrogen-deuterium solvent exchange is so fast in the mutant protein that it is complete before NMR data collection can occur. The former works to reduce the yield of oxyferrous state upon mixture of the reduced heme protein with oxygen, while the latter shortens the lifetime of the oxyferrous state after formation by speeding oxidation during transient pentacoordination subsequent to oxygen binding (see Figure 1D).

Here we demonstrate that changes in protein dynamic motions affect both the rate of water penetration into and hydration of the hydrophobic core of HP-7 by introducing a simple, nonperturbative connecting loop between the two helices which comprise the heme-binding active site. There is a precedent for protein stabilization by fashioning a single-chain protein from a self-assembling helical bundle – when Degrado and coworkers transformed a nonfunctional homotetrameric diporphyrin protein (17) that they had designed into a single-chain tetraporphyrin protein, the overall structure became so stabilized that a number of destabilizing mutations had to me made in order to make the protein flexible enough that it could bind cofactors under non-denaturing conditions (18). Other single-chain four helix heme-binding bundles have been designed (19,20), including one demonstrated to bind oxygen (21), but a detailed examination of dynamic water penetration and oxyferrous state lifetime has not to date been reported.

This minimal modification extends the oxyferrous state lifetime without significantly affecting gaseous ligand binding –the first direct evidence that dynamically-driven water penetration is a rate-limiting step in the oxidation of these complexes. This loop reduces water penetration by limiting the number of possible open conformations, greatly reducing the size of the protein conformational ensemble and consequently the population of open states competent for solvent penetration.

## Materials and Methods

### Chemicals

Hemin was purchased from Fluka (Buchs, Switzerland). Molecular oxygen (O_2_) (99.98% purity), carbon monoxide (CO) (99.9%), and molecular nitrogen (N_2_) (99.99%) gases were from Matheson Gas (Basking Ridge, NJ). N_2_ and CO were scrubbed of residual O_2_ by passage through two bubblers filled with a reduced vanadium sulfate solution followed by another filled with water (22). PD-10 desalting columns were from GE Healthcare (Port Washington, NY). All other solvents and reagents were from either VWR or Sigma.

### General biochemistry

FHFH was expressed and purified as described for other heme protein maquettes earlier (23). Optical spectra were collected with a Hewlett-Packard (New York, NY) 8452A Diode array spectrophotometer running the Olis (Bogart, GA) SpectralWorks software and equipped with a Quantum Northwest (Liberty Lake, WA) Peltier temperature controller. Each binding, reduction potential, and stopped-flow experiment was performed at least three times, and reported errors are standard deviations from the mean. All experiments were performed at 20° C in 250mM Boric Acid, 100mM KCl pH 9.0 unless otherwise specified. Hemin stock solutions of approximately 0.5-1.0 mg/ml were prepared in DMSO and used within four hours. Stock solution concentrations were determined using the pyridine hemochrome assay (24).

Holoprotein complexes, containing 1.0 equivalent per homodimer, were prepared as before (5) by five consecutive additions of 0.2 equivalents of hemin in DMSO with at least ten minutes between additions. Proteins were then purified from any unbound heme using PD10 desalting columns. Holoprotein solution concentrations were determined using the experimentally derived Soret extinction coefficients of ε_414_ = 129 mM^-1^cm^-1^ for HP7-H7F and 122 mM^-1^cm^-1^ for FHFH.

### Circular dichroism spectropolarimetry

CD spectra were recorded on on 15 µM protein a JASCO J-810 sepectropolarimeter in a 1 mm path length quartz cell, using a bandwidth of 1nm and a scan speed of 50nm/min. Background-corrected spectra were collected between 190nm and 250nm. Spectra were collected as ellipticity(θ_obs_, mdeg), and the mean residue ellipticities ([θ],deg•cm^2^•dmol^-1^) were calculated as

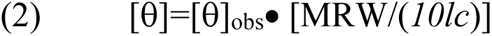

Where MRW is the mean residue molecular weight, *c* is the sample concentration in mg/ml, and *l* is the light-path length of the cell in cm. Protein secondary structure percentages were calculated with the program K2d (25) using data points ranging from 200nm to 240nm.

### Solution molecular weight determination

Oligomerization states were determined using size exclusion chromatography on a Biologic DuoFlow FPLC pump (Biorad Inc., Hercules, CA) equipped with a Quadtek detector and a Bio-Rad Bioselect 125-5 300×7.6 mm column. Protein was eluted with 50 mM NaH2PO4, 10 mM NaCl pH 7.5 at a 0.5 ml/min flow rate. Protein elution was monitored at 280 nm. The column was standardized using Biorad gel filtration standards chicken ovalbumin, α-lactalbumin, and conalbumin (14-75 kDa).

### Heme affinity measurements

For oxidized binding experiments approximately 0.1 molar equivalent aliquots of hemin solution were added via gastight syringe to a stirring 4 ml solution of 2-3 μM protein, with a ten minute equilibration delay between additions. Heme binding was monitored by loss of the absorption at 385 nm due to free hemin, and the concomitant appearance of a sharp Soret band at 412 nm, corresponding to heme bound to the protein via bis-histidine axial coordination. The number of heme binding sites was quantified for each protein from plots of the Soret maximum at 412 nm vs the number of equivalents added.

Reduced binding experiments were performed on solutions of ∼3 μM protein as described previously (5). Briefly, hemin was added in approximately 0.1 molar equivalent aliquots to anaerobic protein solution kept at a potential less than −450 mV using periodic additions of sodium dithionite. K_d_ values were obtained from plots of the Soret band absorbance measured at 436 nm vs. the concentration of hemin added and fit with the tight binding equation:

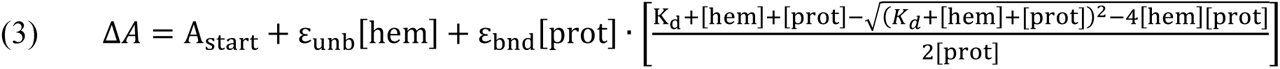

Where ε_unb_ is the molar absorption coefficient of unbound hemin at that wavelength, ε_bnd_ is the additional absorbance of bound hemin at that wavelength, [hem] is the total hemin concentration, [prot] is the total protein concentration and K_d_ is the dissociation constant for the reduced hemin.

### Reduction potential determination

Spectroelectrochemical redox titrations were performed as described previously (5) using 10-20 *μ*M solutions of hemoprotein containing >100 μM of the corresponding apoprotein in order to eliminate the possibility of heme dissociation upon reduction. Reported reduction potentials are referenced to a standard hydrogen electrode. All spectroelectrochemical titrations were performed anaerobically using *μ*L additions of freshly prepared sodium dithionite to adjust the solution potential to more negative values and potassium ferricyanide to more positive values. Redox mediators were as used previously (26). Titrations were analyzed by monitoring the Q band absorbance at 559 nm as the heme protein was reduced or oxidized and data fit with the Nernst equation using an *n*-value of 1.0:

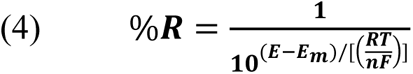

where %R is the fraction of reduced heme, E is the solution potential, E_m_ is the reduction midpoint potential, and n is the number of electrons.

### Flash photolysis analysis of distal histidine association

Rate constants for CO and histidine binding to the pentacoordinate state were determined using laser flash photolysis. Transient absorbance data were collected on an Edinburgh Instruments LPK920 laser flash photolysis spectrometer equipped with an Opotek Vibrant 355II tunable laser source. Excitation at 532 nm was used to excite the preformed carbonmonoxyferrous complex causing the detachment of the ligand CO and the rates and magnitudes of CO and histidine association were determined by analyzing multi-exponential rebinding traces taken as a function of CO concentration using the method of Hargrove (27):

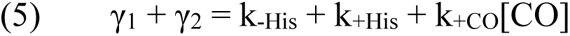

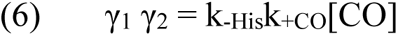

Where γ _1_ and γ _2_ are the fitted first and second CO-dependent exponential rates and the kinetic constants are defined as in Equation 7. Protein concentrations were 20-25 μM and carbonmonoxyferrous complexes were prepared by reducing solutions of the holoproteins with an excess of dithionite under an atmosphere containing 10-100% CO mixed with N_2_.

### Stopped Flow analysis of gaseous ligand binding

Binding kinetics of O_2_ and CO with ferrous protein-heme complexes were determined as before (4) using a Biologic (Lyon, France) SFM 300 stopped flow mixer equipped with a custom-built Olis RSM 1000 spectrometer for multiwavelength detection. Binding was followed spectroscopically over gas concentrations from 2% to 50% saturation at 15°C. Air- or gas-saturated buffer was mixed with N_2_-saturated buffer in the first mix, and then this mixture was combined in a second mix with an N_2_-saturated ferrous heme protein solution. Protein concentrations were 20 μM and ferrous complexes were prepared by carefully titrating anaerobic solutions of the holoproteins with a slight excess of sodium dithionite as observed by visible spectroscopy followed by anaerobic canular transfer to the stopped-flow loading syringe. Binding kinetic data were fit with Eqn. 7, which assumes that O_2_ binding rate, k_+O2_, is much greater than the sum of the distal histidine association and dissociation rates (28):

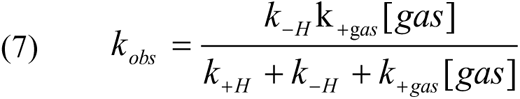

Where k_obs_ is the fitted single exponential binding rate, k_+H_ and k_-H_ are the distal histidine-ferrous heme iron association and dissociation rates, and k_+gas_ is the gaseous ligand association rate constant. At high [gas], k_obs_ = k_-H_.

### Kinetic analysis of CO dissociation

The dark dissociation rate of the CO complex was determined using the ferricyanide trapping method of Moffet *et al* (27): briefly, anaerobic solutions of carbonmonoxyferrous protein complex were mixed with varying concentrations of potassium ferricyanide and the rates of the linked reactions of CO dissociation followed by ferricyanide heme oxidation were followed by monitoring the disappearance of the carbonmonoxyferrous Soret peak at 421 nm. The rates of oxidation *vs.* the concentration of ferricyanide give the inverse CO dissociation rate as given by:

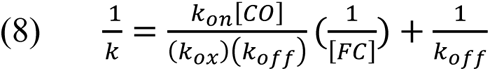

### Kinetic analysis of O_2_ dissociation

The dissociation rate of the O_2_ complex was determined using the CO displacement method of Gardner et al (29). Briefly, the O_2_ complex was formed as above by mixing reduced protein and O_2_-saturated buffer in a stopped-flow spectrophotometer followed by a second mix 100 ms later with a two-fold larger volume of CO-containing buffer. The rate of O_2_ replacement by CO was collected using final CO concentrations ranging from 200-600 µM. At high CO concentration, the observed rate of replacement, r_obs_ is given by:

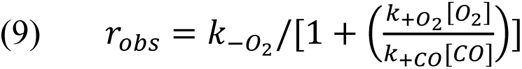

When *k***_+_**_co_[*CO*] ≫*k*_+O2_[O_2_], the observed replacement rate constant is directly equal to O_2_ dissociation rate constant: r_obs_≈ *k*_−*O*2_.

### Oxyferrous state lifetimes

As before (4), the lifetimes of O_2_-bound ferrous protein-heme complexes were determined spectroscopically in stopped-flow mixing experiments using final O_2_ concentrations from 16.67% to 66.67% saturation at 16° C. Protein concentrations were 10 μM with 1 μM hemin added to ensure full complexation. Ferrous complexes were prepared by carefully titrating anaerobic solutions with a slight excess of sodium dithionite as observed by visible spectroscopy. Ferrous samples were anaerobically transferred to the stopped-flow loading syringe by canula. Kinetic data were fit with Eqn. 1.

### Nuclear magnetic resonance

All NMR experiments were performed at 20°C using a Varian Inova spectrometer operating at a 600MHz and equipped with a triple resonance cryogenic probe capable of applying pulse field gradients in the z-direction. Chemical shifts are referenced to water at 4.77 ppm for ^1^H. Data were processed using the program NMRPipe (30) and analyzed using Sparky (31).

Sensitivity enhanced ^1^H-^15^N heteronuclear single quantum coherence (HSQC) spectra (32) were collected to assess structural specificity (23) on 50-100 μM holo- and apoprotein samples with sweep widths of 10,000 Hz for ^1^H and 2000 Hz for ^15^N utilizing GARP decoupling of ^15^N during ^1^H acquisition.

In order to determine the pK_a_ of histidine side chains, the ^1^H-^15^N multiple-bond correlation signals originating in the imidazole side chains were collected as before (5) using a non-sensitivity enhanced HSQC pulse sequence in which the INEPT (insensitive nuclei enhanced by polarization transfer) periods were set to 1/^1^J_NH_, where ^1^J_NH_ is the one-bond ^1^H-^15^N coupling constant, in order to attenuate backbone amide signals (33). Spectral widths were 8000 Hz for ^1^H and 12156 Hz for ^15^N.

^15^N-labelled apoprotein samples were 200 µM in 25mM KaD_2_PO_4_ D_2_O buffer with the pD (pH meter reading + 0.4 pH units) initially fixed at 5.0. After each spectrum was taken the pD was adjusted upwards by the addition of small amounts of KOD dissolved in D_2_O. The observed ^15^N chemical shifts were fit with the Henderson-Hasselbach equation assuming fast chemical exchange between protonation states:

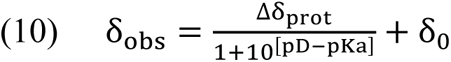

where δ_0_ is the neutral chemical shift of ^15^N, Δδ_prot_ is the change in chemical shift due to protonation, and pK_a_ is the fitted acid dissociation constant.

### Hydrogen-Deuterium exchange

To compare the rates of backbone hydrogen exchange (34), ^15^N-labelled holo- and apoprotein samples of HP7-H7F and FHFH in 25 mM potassium phosphate buffer pH 6.5 were passed through a PD-10 solvent exchange column pre-equilibrated in 25 mM KD_2_PO_4_ D_2_O buffer pD 6.5 (pH meter reading + 0.4 pH units), immediately placed in a 5mm NMR tube, and equilibrated at 20°C for five minutes in the sample compartment of the NMR spectrometer. Final protein concentrations were approximately 300 μM. The fraction of remaining amide protons was assessed as a function of time by collecting one dimensional ^15^N-HSQC, or isotope-selective, ^1^H spectra.

### Deep UV resonance raman (dUVRR) spectroscopy

The dUVRR spectra of the FHFH and HP-7-H7F were acquired as described previously (35) using a custom-built instrument (36). Incident laser power at the sample was attenuated to 500 µW to avoid protein degradation, and spectra were monitored for degradation over time using the aromatic ring modes. Spectral calibration was carried out using a standard cyclohexane spectrum (37). All dUVRR spectral preprocessing was carried out in MATLAB (7.1, MathWorks, Natick, MA) using cosmic ray and water band removal methods described previously (38). A nonlinear least-squares algorithm was used to fit the amide and aromatic bands to mixed Gaussian/Lorentzian peaks, which approximate the Voigt line shape as described previously (39).

As standard reductants such as dithionite have significant molar absorptivity coefficients in the deep-UV, spectra of the reduced forms of each heme protein were acquired using titanium(III) citrate as the reductant. Concentrated solutions of titanium(III) citrate were generated by adding 5 mL of an Ar purged buffered solution of 0.2 M sodium citrate (pH 7) to approximately 75 mg of TiCl_3_ to produce titanium(III) citrate in an Ar purged crimp top vial. H4 and HP7 protein samples (4.5 µM and 5.3 µM) were prepared in 25 mM borate, 10 mM KCl, 50 mM NaClO_4_ buffer (pH 9.5) and purged with Ar in similar vials. Air exposed samples were confirmed for reduction and reoxidation by UV-Vis spectroscopy. 5 µL aliquots of titanium(III) citrate solution were added to 1 mL of each protein sample until the absorption band at 560 nm reached a maximum intensity samples remained fully reduced for 15-20 minutes, which defined the maximal time frame for individual sample collection in the dUVRR sample flow cell.

## Results

### Protein design

Water penetration into the hydrophobic core of a globular protein like HP7-H7F is driven by large-scale or even global dynamic protein motion (34). In order to restrict these global motions, we surmised that a simple glycine-rich connecting loop similar to that we previously used to create the light-activated electron transfer dyad protein HHHF (40) could reduce water penetration by limiting the number of possible open conformations, greatly reducing the size of the protein conformational ensemble and consequently the population of open states competent for solvent penetration. Thus we modified the homodimeric “candelabra” structure (23) by connecting the second helix of one monomer of HP7-H7F to the first helix of the other, resulting in the single chain protein FHFH (Figure 1A-C).

FHFH displays visible absorbance spectra indicative of bis-histidine coordination in both oxidation states (Figure 3A) with Soret and Q-band peak maxima identical to those of heme bound HP7-H7F. It is important to confirm the helical bundle structure of the protein, as some designed helical bundle proteins have been found to form domain-swapped higher oligomeric states (41). Both apo- and holoprotein forms of FHFH are monomeric as analyzed by size exclusion chromatography (Supplemental Figure 1). Furthermore, while the apoprotein appears to be helical as detected by circular dichroism (Supplemental Figure 2), it has ^15^N-HSQC spectra indicative of molten globular structure. The addition of a single heme cofactor changes the HSQC spectra to a level of chemical shift dispersion indicative of a partial phase transition in which the two helices which ligate the heme become native-like and the two unliganded helices remain molten globular, consistent with what we have previously observed for HP7 (Supplemental Figure 3) (23).

### Core hydration

*Interior group pKa’s:* Several different lines of evidence suggest that the introduction of this loop significantly reduces the hydration of the protein interior: First, the pK_a_ of the buried ligand histidine side chains in the apoprotein is 1.4 pH units lower in FHFH than in HP7-H7F (see Figure 2A and supplementary Figure 4), corresponding to an increase of 1.9 kcal/mol in the energy required for protonation. While there are many local structural interactions which can shift side chain pK_a_s in either direction, the most common cause of pK_a_ shifts which destabilize the charged state of side chains in proteins is a decrease in the local dielectric (42).

**Figure 2.**
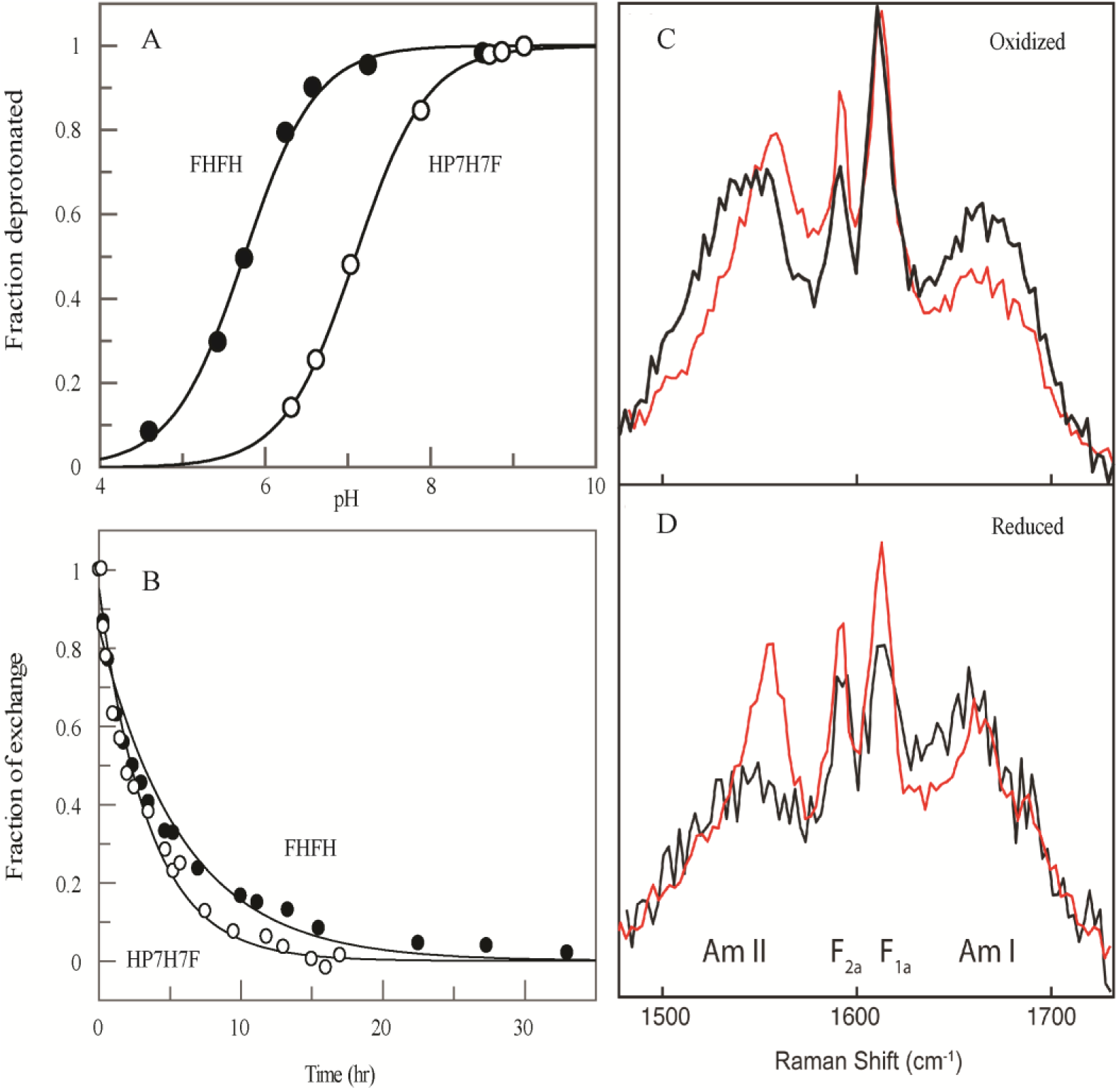
Experimental demonstration of reduced hydration in the core of FHFH. (A) NMR determination of the pK_a_ values of the histidine ligands in apo-HP7-H7F (○) and FHFH (●). Lines drawn are fits of the ^15^N(ε2) chemical shifts with equation 10 having pK_a_ values of 7.3 and 5.9 respectively. (B) Hydrogen-deuterium exchange in the ferric heme complexes of HP7-H7F (○) and FHFH (●). Lines drawn are exponential fits to the integrated data with exchanges rates of 0.17 s-1 and 0.29 s-1 respectively. (C and D) Deep-UV resonance Raman spectra of the oxidized (C) and reduced (D) forms of the HP7-H7F (black) and HFHF (red) are shown for the spectral region containing the backbone Amide I and II mode as well as two hydration-sensitive ring related aromatic modes.

### Hydrogen/Deuterium exchange

A more rigorous method of comparing interior hydration is that of hydrogen/deuterium exchange (43). In whole-protein experiments, the bulk of backbone amide residues exchange in a few seconds at neutral pH and 25°C. These are the residues either directly exposed to or only minimally protected from solvent. Exchange of core residues, however, is typically greatly slowed – core proton exchange requires an open conformation which exposes these backbone amides to solvent. Such conformations are much higher in energy than lowest energy or ‘Native-state’ conformations. Exchange rates for these residues are thus the sum of the fractional Böltzmann populations of all of the open states competent for hydrogen exchange multiplied by the exchanges rates of these conformations.

We have previously shown that the core residues of ferric HP7-H7F exchange on a minutes time scale, too rapidly to collect two dimensional data on individual residue protection factors but slowly enough that 1-dimensional ^15^N-selective ^1^H NMR data on the exchange of the entire amide proton population can be quantified (4). A similar one-dimensional analysis of ferric FHFH demonstrates that core residue hydrogen exchange is almost two-fold slower (Figure 2B). Thus the addition of the loops restricts water access to the protein core, either by making a subset of these open conformations impossible to form or too high in energy to populate, effectively reducing the internal water concentration.

### Deep UV Resonance Raman

A complimentary method for examining the hydration of protein interiors is dUVRR. This vibrational technique utilizes resonance enhancement of both the peptide backbone and aromatic residue Raman modes to provide information about the ensemble hydration status of the backbone (amide I and II modes) as well as more regiospecific information via aromatic residue environment-derived changes. Here we focus in particular on the phenyalanine modes F_1a_ and F_2a,_ as this is the only aromatic residue type in the protein core – the sequences contain no tyrosine side chains and the tryptophan rings are purposely placed only at the solvent-exposed C-position termini of each helix for the purpose of protein concentration determination (26). Both the F_1a_ and F_2a_ mode intensities, as well as the intensity ratio of the amide I and amide II modes, have been shown in other proteins and model systems to change as a function of regional side chain dehydration (44–46).

As Figure 2C and 2D demonstrates, the magnitudes of the phenylalanine (F_1a_ and F_2a_) and amide II mode intensities in both the ferric and ferrous heme complexes of the two proteins are markedly different. These differences are much larger than we have observed for the association of the helical pH low insertion peptide (pHLIP) with membranes and are closer to those we see for membrane insertion of pHLIP (47). These data clearly demonstrate that the protein interiors of both the ferric holo and ferrous holo states of FHFH are dehydrated in comparison to the same states of HP7-H7F, further evidence that the addition of the loop indeed does restrict the size of the conformational ensemble.

### Ferric and ferrous heme binding

Given the change in core hydration without a significant change in protein structure, it is important to determine the effect the introduction of this loop has on the binding of the heme cofactor. As has been observed for HP7-H7F (5), ferric heme binding to FHFH is too tight to reliably extract a dissociation constant. This enabled us to perform endpoint titrations, confirming that FHFH binds a single heme cofactor as well as calculate the extinction coefficient of the oxidized complex (Figure 3A).

**Figure 3.**
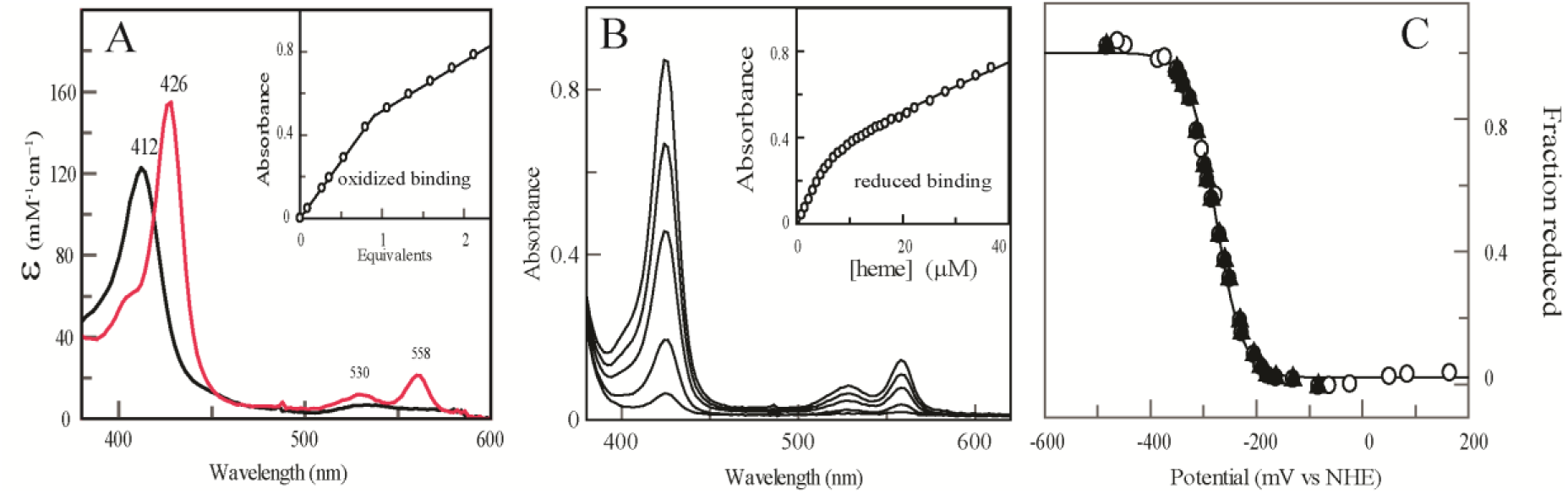
The thermodynamics of heme binding to FHFH. (A) Oxidized (black) and reduced (red) spectra and oxidized heme titration (inset) of heme-bound FHFH. Extinction coefficients were calculated using the intercepts from the end point titrations. (B) Reduced heme binding to 3.0 *μ*M FHFH. Some spectra have been omitted for the sake of clarity. (Inset) Equilibrium binding isotherm derived from the titration in panel B. The line drawn is a fit with Eqn. 3 using a *K*_d,red_ of 3.4 µM. (C) Equilibrium potentiometric determination of the electron affinity of FHFH. Line drawn is a fit with eq 3 with an E_mid_ of −277 mV vs NHE.

Ferrous heme binding, however, is weak enough that binding energies can be quantitated under reducing conditions (Figure 3B). Ferrous heme binding is more than five-fold weaker in FHFH than HP7-H7F (Table I). The reduction potential of the bound heme in FHFH is 17 mV lower than that of HP7-H7F (Figure 3A), and in combination with the free heme reduction potential of −63 mV vs. NHE enables the calculation of the ferric heme binding affinity using a thermodynamic cycle (48). The calculated ferric heme affinity, 900 pM, is itself three-fold weaker than that of HP7-H7F.

**Table I.** Heme binding parameters for two artificial heme proteins.

| Protein | $K_{d-red}^*$ (μM) | $E_{mid}^\dagger$ (mV) | $K_{d-ox}^\ddagger$ (nM) | $k_{+his}$ ($s^{-1}$) | $k_{-his}$ ($s^{-1}$) | $K_{A,his}^\S$ | $K_{d_{pen}}^\parallel$ (μM) | $pK_a^\P$ |
| --- | --- | --- | --- | --- | --- | --- | --- | --- |
| HP7-H7F | $0.6 \pm 0.2$ | $-260 \pm 6$ | $0.3 \pm 0.1$ | $160 \pm 20$ | $5.6 \pm 0.3$ | $29 \pm 5$ | $18 \pm 9$ | $7.3 \pm 0.2$ |
| FHFH | $3.4 \pm 0.5$ | $-277 \pm 1$ | $0.9 \pm 0.1$ | $210 \pm 40$ | $24 \pm 1$ | $9 \pm 2$ | $34 \pm 8$ | $5.9 \pm 0.2$ |

\* hexacoordinate dissociation constant in the reduced state

### Distal histidine affinity

A possible reason for the observed decrease in ferrous heme affinity in FHFH is a decrease in distal histidine affinity caused by the decreased dielectric in the protein core. We have previously shown that the overall heme binding affinity to hexacoordinate hemoglobins is (5):

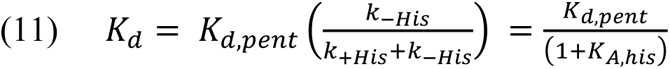

Where K_d,pent_ is the dissociation constant for the pentacoordinate heme complex and K_A,his_ is the distal histidine association constant – the ratio of the distal histidine on-(k_+His_) and off-rates (k_-His_). k_-His_ were determined using CO binding: as the protein is primarily in the hexacoordinate bis-histidine ligation state in solution (Table 2) and the rate-limiting step for CO binding at high ligand concentrations is the detachment of the distal histidine from the heme iron (see eq 6).(5) Figure 4A depicts the concentration dependence of the CO binding rate to both proteins. Comparison of the asymptotic limits demonstrates that the histidine off-rate is four-fold slower in FHFH than HP7-H7F while k_+His_ rates, determined using the laser flash method of Hargrove (Figure 4C) (49), were within error of each other (Table I). Thus the decreased binding energy indeed seems to primarily result from decreased distal histidine affinity, which is itself likely a result of the increase in the energy required to move polar glutamic acid side chains into a lower dielectric environment.

**Figure 4.**
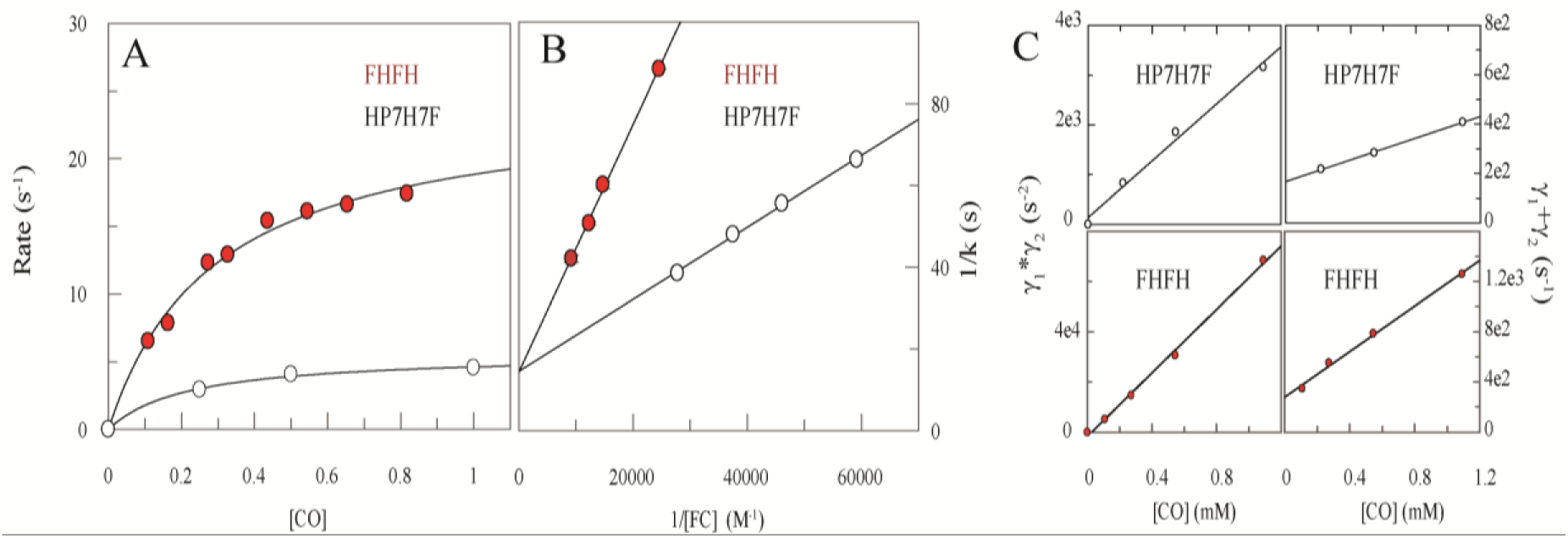
Kinetic analysis of CO and histidine binding to HP7-H7F (white circles) and FHFH (red circles). (A) Stopped-flow analysis of the rates of binding of CO to reduced heme proteins as a function of ligand concentration. Lines are fits with Eqn. 7. (B) Ferricyanide trapping analysis of CO release from FHFH and HP7H7F. Double-reciprocal plots of the rates of oxidation vs the concentration of ferricyanide extrapolated to infinite ferricyanide give the CO dissociation rate as shown in eq 7. (C) Laser flash kinetic analysis of histidine and CO rebinding. Flash data were fit with three exponentials, the first of which represents a CO concentration-independent relaxation process (see ref. 5). Replots of the products (left) and sums (right) of the second two exponentials. Fits with Eqns. 5 and 6 give the k+His and k+CO.

### Gaseous ligand binding

Figures 4B and 4C depict the experimental determination of the rates of CO binding (k_+CO_) and dissociation (k_-CO_) to both proteins. Both rates and the derived pentacoordinate binding constants, K_d,CO_pent_, are within error of each other (Table II). One significant difference is seen in the ferricyanide trapping analysis of CO release (Figure 4B) – although the intercepts and thus the dissociation rates are equivalent, the slopes of the replots are significantly different, perhaps another result of restricted access to the heme in FHFH. The true binding constants involve competition between gaseous ligand binding and distal histidine coordination (28):

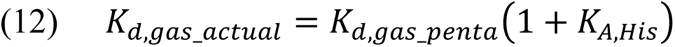

Therefore the actual binding CO dissociation constants differ by a factor of four.

Similar behavior is observed with oxygen: Figure 5 depicts the stopped-flow analysis of the O_2_ on- and off-rates of both proteins. Both pairs of rates are identical within experimental error (Table II), but faster than what is observed for CO. Likewise, both pentacoordinate binding constants, K_d,O2_pent_, are within error of each other, and the difference in K_A,His_ is a factor of three. Thus loop addition has no measurable effect on the binding of either ligand.

**Figure 5.**
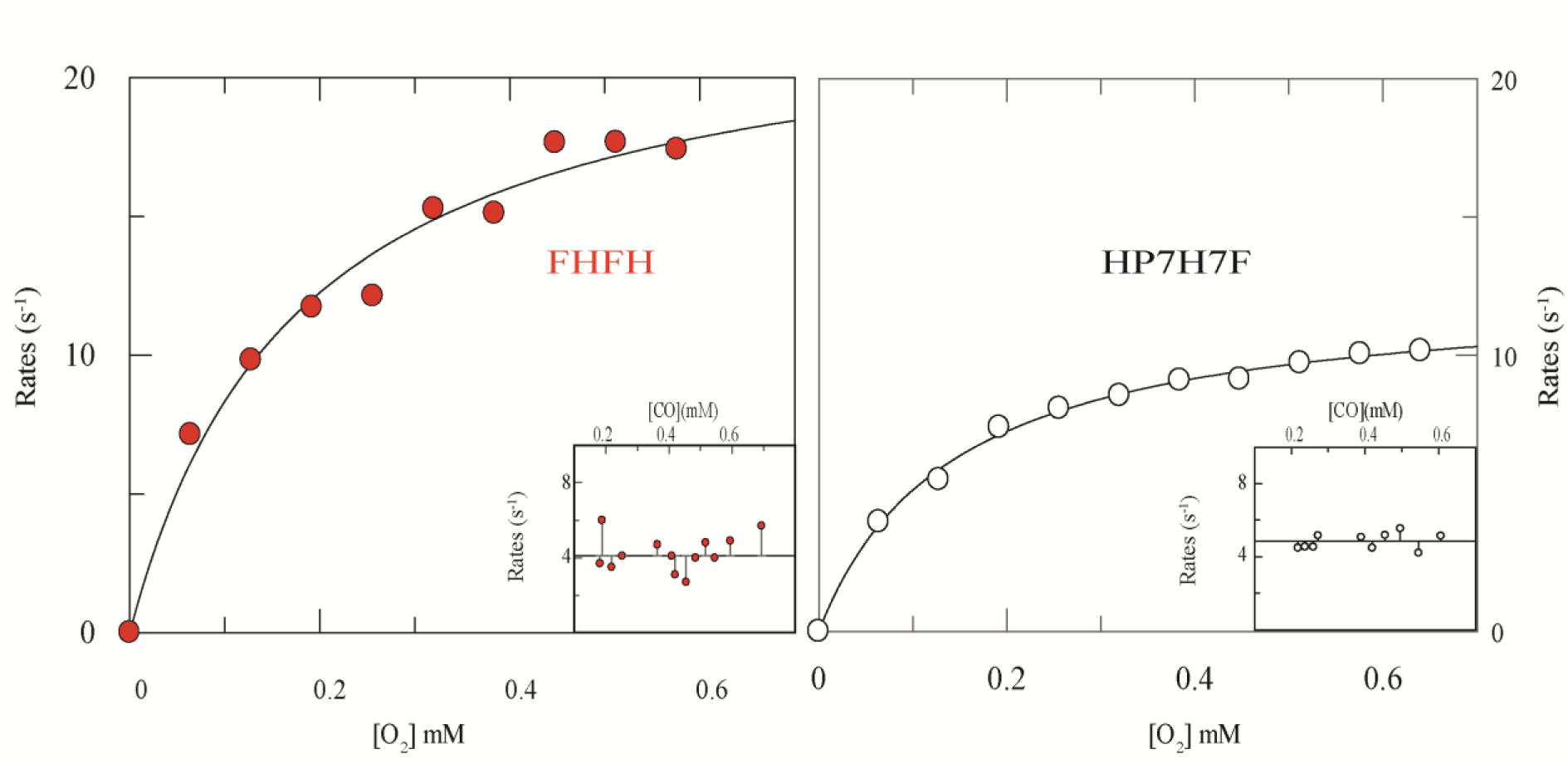
Kinetic analysis of O2 binding to FHFH (red circles) and HP7-H7F (white circles). On-rates as a function of oxygen concentration. Reduced protein−heme complex (20 μM), prepared by careful titration with a slight excess of dithionite, was mixed with oxygen in a stopped-flow apparatus, and the rate of oxyferrous state formation was followed spectroscopically. Lines shown are fits to the data with eq 1. (Insets) Off rates as a function of carbon monoxide concentration. Lines are the average rates of oxygen displacement.

**Table II.**
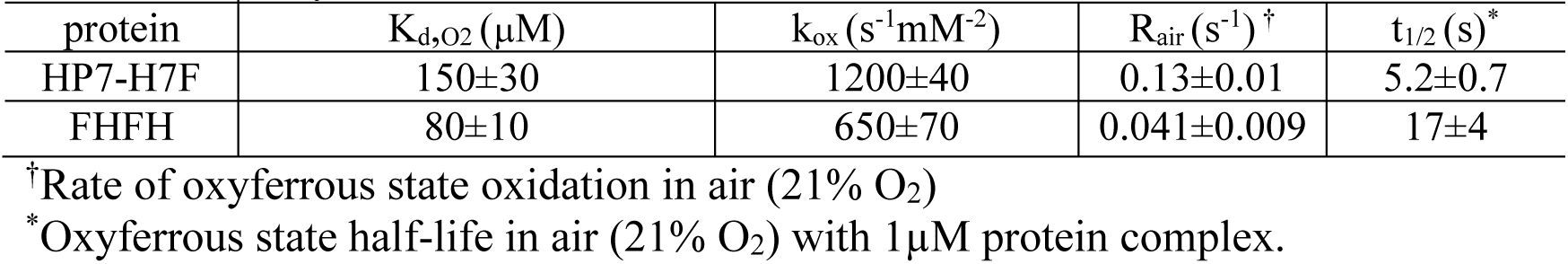
Gaseous ligand on- and off-rates and equilibrium constants at pH 9.

### Oxyferrous state lifetime

Under standard conditions (1 µM heme-protein complex, 21% O_2_) FHFH has an oxyferrous state lifetime more than threefold longer than HP7-H7F (Figure 6A). Figures 6B depict the O_2_ dependence of the lifetime. Fitted O_2_ dissociation constants agree well with those determined using the ratio of on and off rates (see Table III). FHFH has an oxidation rate two-fold slower than that of HP7-H7F.

**Figure 6.**
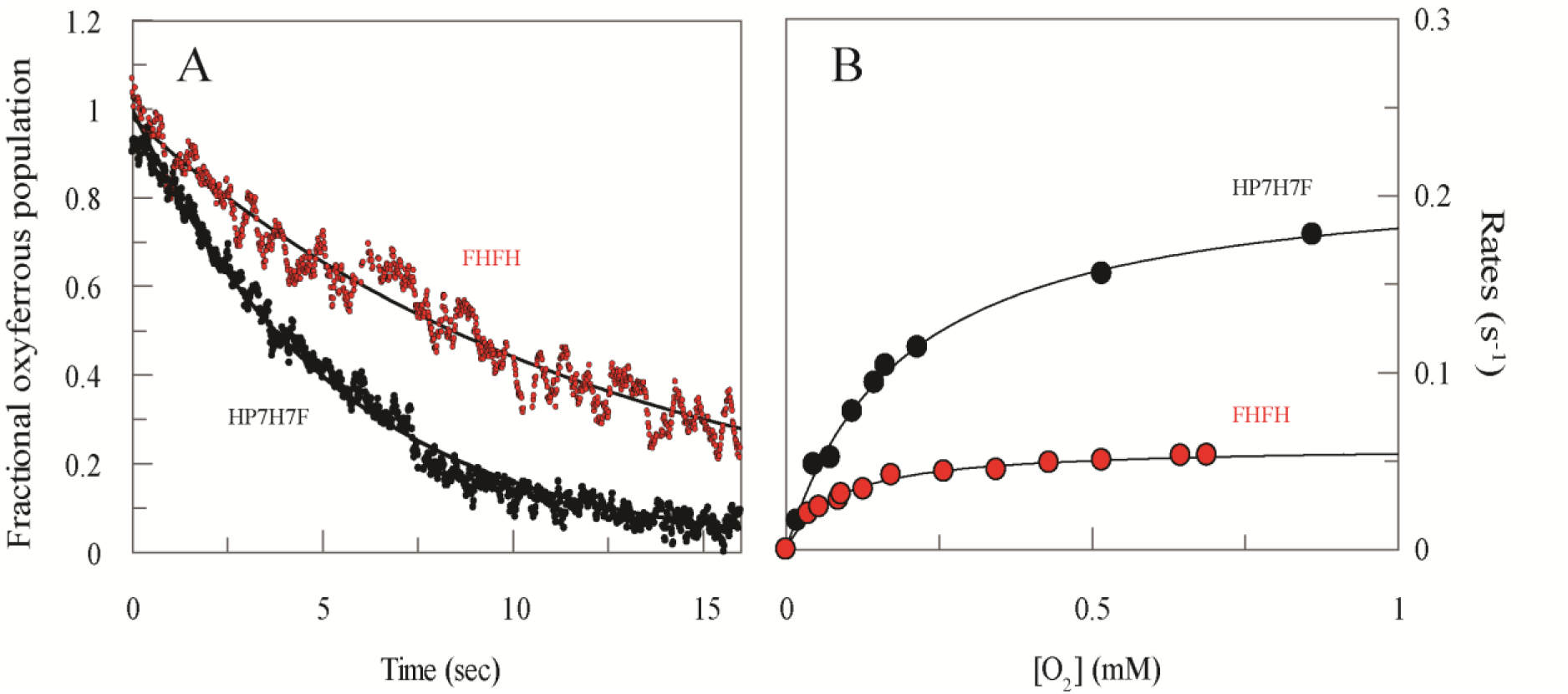
Rates of oxyferrous state breakdown in FHFH (red circles) and HP7-H7F (black circles). Reduced protein−heme complex (1 μM), prepared by careful titration with a slight excess of dithionite, was mixed with oxygenated buffer and the rate of protein oxidation followed spectroscopically. (A) Direct comparison of the breakdown of 1 µM oxyferrous state in air (21% saturated O2). Lines are fits with a single exponential function with rates of 0.041 s-1 and 0.13 s-1. (B) Oxygen concentration dependence of the oxyferrous state breakdown of FHFH (red circles) and HP7-H7F (black circles) respectively.

**Table III.**
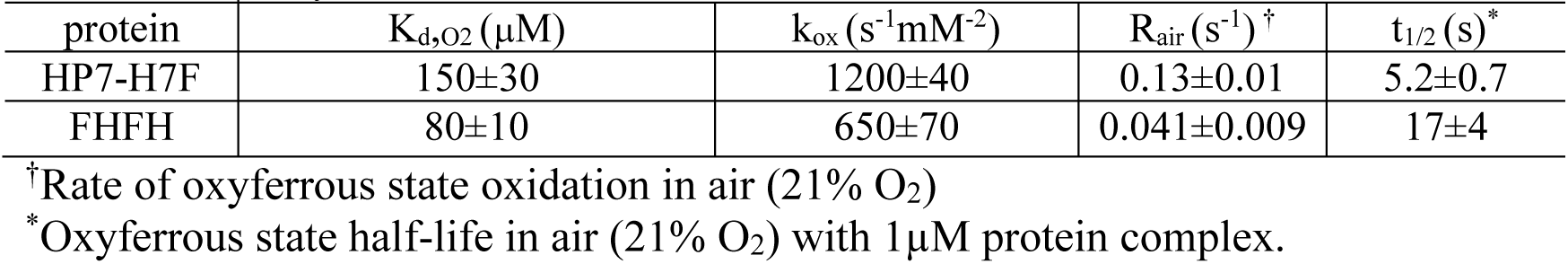
Oxyferrous state lifetimes.

## Discussion

The identical pentacoordinate binding constants of both ligands to each protein in concert with the indistinguishable spectra of the bound states suggests that the bound ligand environments are identical in both proteins. Therefore the addition of the connecting loop significantly reduces water access, and thus the internal dielectric, of the protein core without significantly altering the heme binding site. As one would expect, this has a small effect on the binding of the heme cofactor but no measurable effect on the binding of uncharged gaseous ligands. It does have a significant effect on protein function – the lifetime of the oxyferrous state increases by a factor of more than three. Two parameters play a role in oxyferrous state lifetimes, the O_2_ binding constant and the second order rate constant for the oxidation of ferrous heme by unbound oxygen.

The slower oxidation rate, which correlates with a decrease in the rate of water penetration, is the principal basis for the increase in oxyferrous state lifetime. The distal histidine dissociation rate, which is also altered by the loop addition, will not affect the lifetime as both proteins have oxygen on and off-rates that are more than an order of magnitude faster than the histidine on-rate. Therefore once oxygen is bound, oxidation of the reduced heme occurs only after oxygen transiently dissociates, allowing the pentacoordinate heme to be oxidized by a molecule of oxygen (50). The increased distal histidine off-rate will in theory result in an increased yield of oxyferrous complex upon mixing ferrous protein with oxygen, but HP7-H7F already quantitatively forms the oxyferrous state within experimental error. Furthermore the lower reduction potential of FHFH should serve to destabilize the oxyferrous state, making oxidation of the heme more favorable. Therefore, it is unlikely to be responsible for the increase in lifetime and we attribute the lengthened lifetime directly to the decreased water penetration into the protein core. To our knowledge, this the first direct demonstration that dynamically-driven water penetration can be a rate-limiting step in the oxidation of protein-heme complexes.

This relationship between protein dynamics, water penetration, and the stability of metalloprotein intermediate states is instructive as to the reasons for the failure of earlier designed heme proteins to stably bind oxygen: early minimalist peptide-protein complexes directly expose the heme to solvent (51,52). The first designed heme proteins large enough to entirely occlude the heme surface were composed of helical pairs, crosslinked by disulfide bond formation between terminal cysteine residues, which tetramerized via hydrophobic sequestration (53–55). However, these proteins are molten globules with the attendant rapid water penetration. The next generation of designed proteins formed using this topology are uniquely structured in the oxidized state (26,56), but the disulfide bonds which hold the fold together have reduction potentials higher than the bound heme cofactors (57). This results in the formation of a molten globular state upon heme reduction and the concomitant reduction of the protein disulfides, and explains why these reduced proteins cannot stably bind oxygen. The candelabra fold, which is the scaffold for HP7, consists of two helix-loop-helix chains which bind two hemes at the interface between the two monomers (23). The effect of this is to add sufficient rigidity to restrict water penetration such that there is measurable amide proton hydrogen exchange protection (4) and stable formation of an oxyferrous state. The increased rigidity imparted by the addition of the connecting loop in FHFH further restricts water penetration and thus further increases the stability and lifetime of the oxyferrous state.

In conclusion, these data are not only instructive as to the engineering requirements for natural oxygen transport and oxygen activation proteins; they make it clear that an important design principle in the construction of functional metalloproteins and metalloenzymes is the restriction of water from the active site.

## Author Contributions

L.Z., M.C.B., A.C.M., K.N.G., J.W.C. and R.L.K designed research; L.Z., M.C.B., A.C.M., and K.N.G., performed research; L.Z., M.C.B., J.W.C. and R.L.K wrote the manuscript.

## Notes

The authors declare no competing financial interests.

## Acknowledgements

The authors would like to thank John Olson, of the department of Biochemistry and Cell Biology, Rice University, for helpful discussion of this manuscript.

---

**Supplemental Figure 1.**
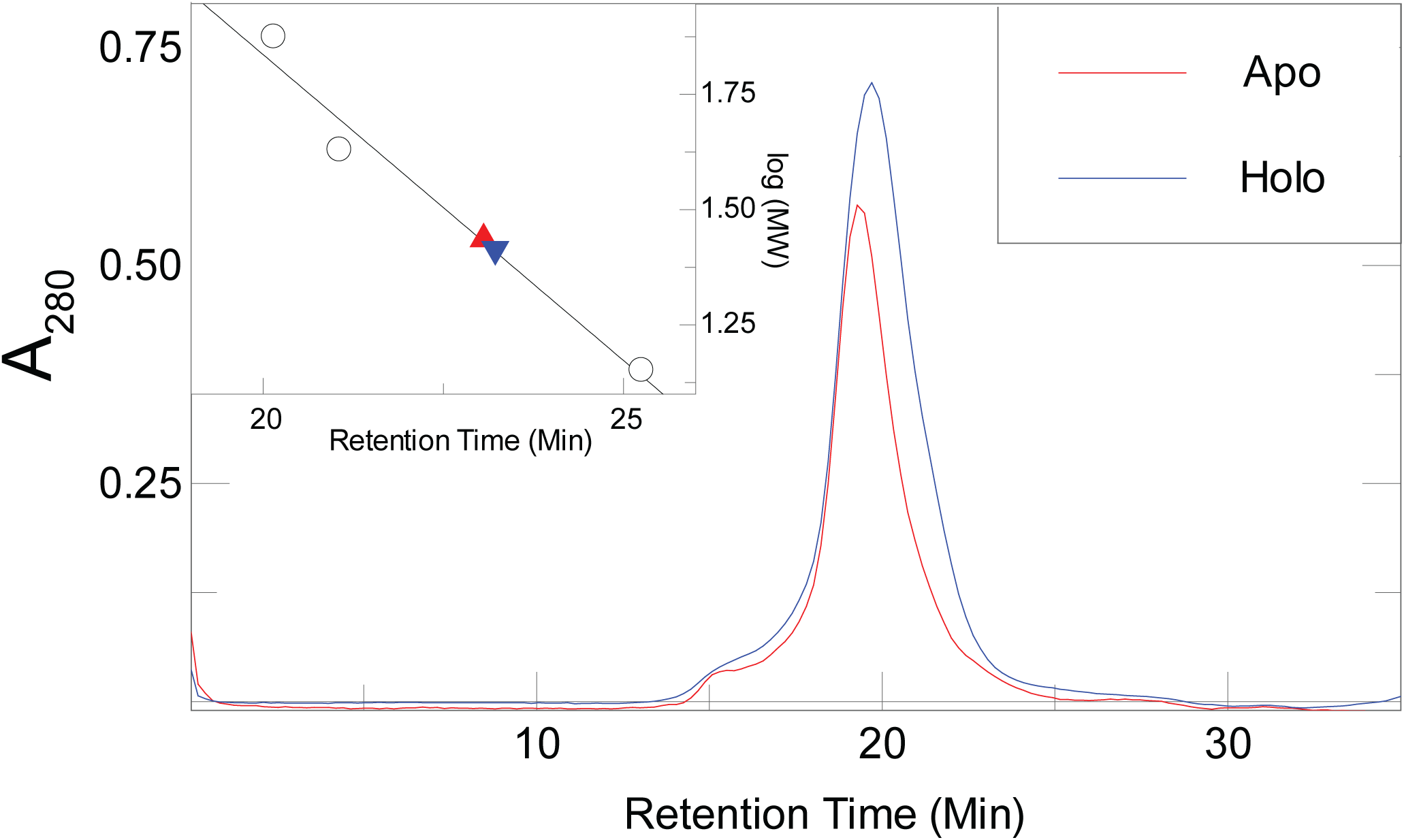
Size exclusion chromotogograms of apo-FHFH (red) and holo-FHFH (blue). Inset: Calibration curve used to estimate the molar mass and solution oligomeric state of each protein: 23.1 (apo) and 23.2 (holo) kDa.

**Supplementary Figure 2.**
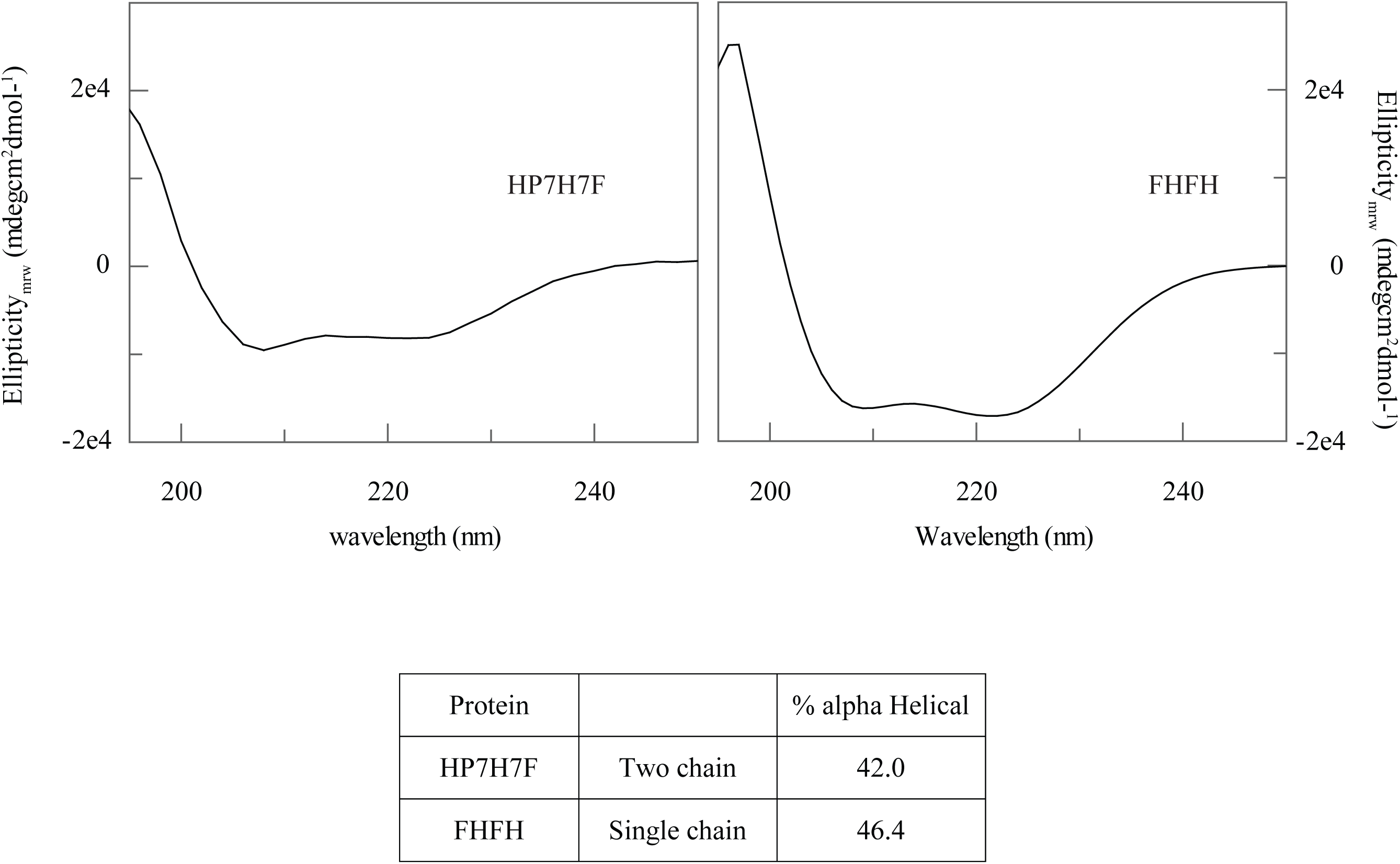
Circular Dichroism spectra

**Supplementary Figure 3.**
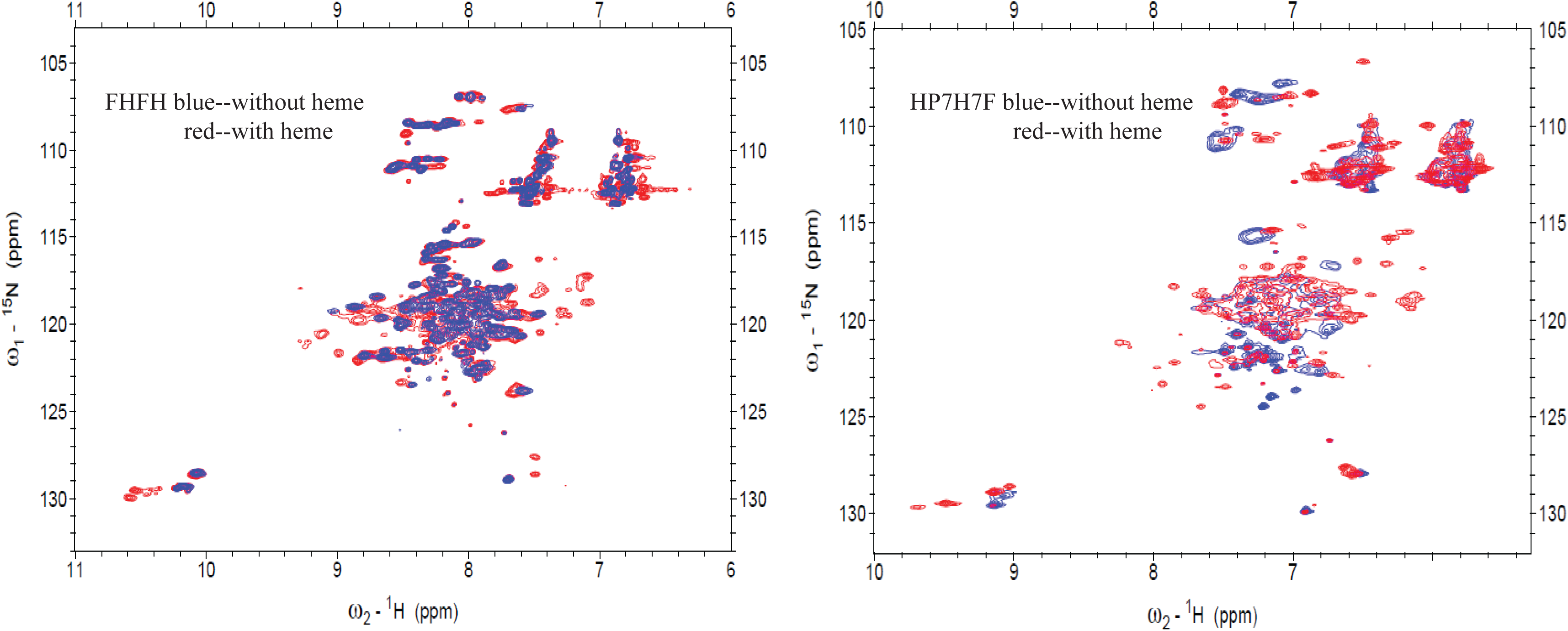
^15^N HSQC specta of apo- and holo-HP7H7F and FHFH

**Supplementary Figure 4.**
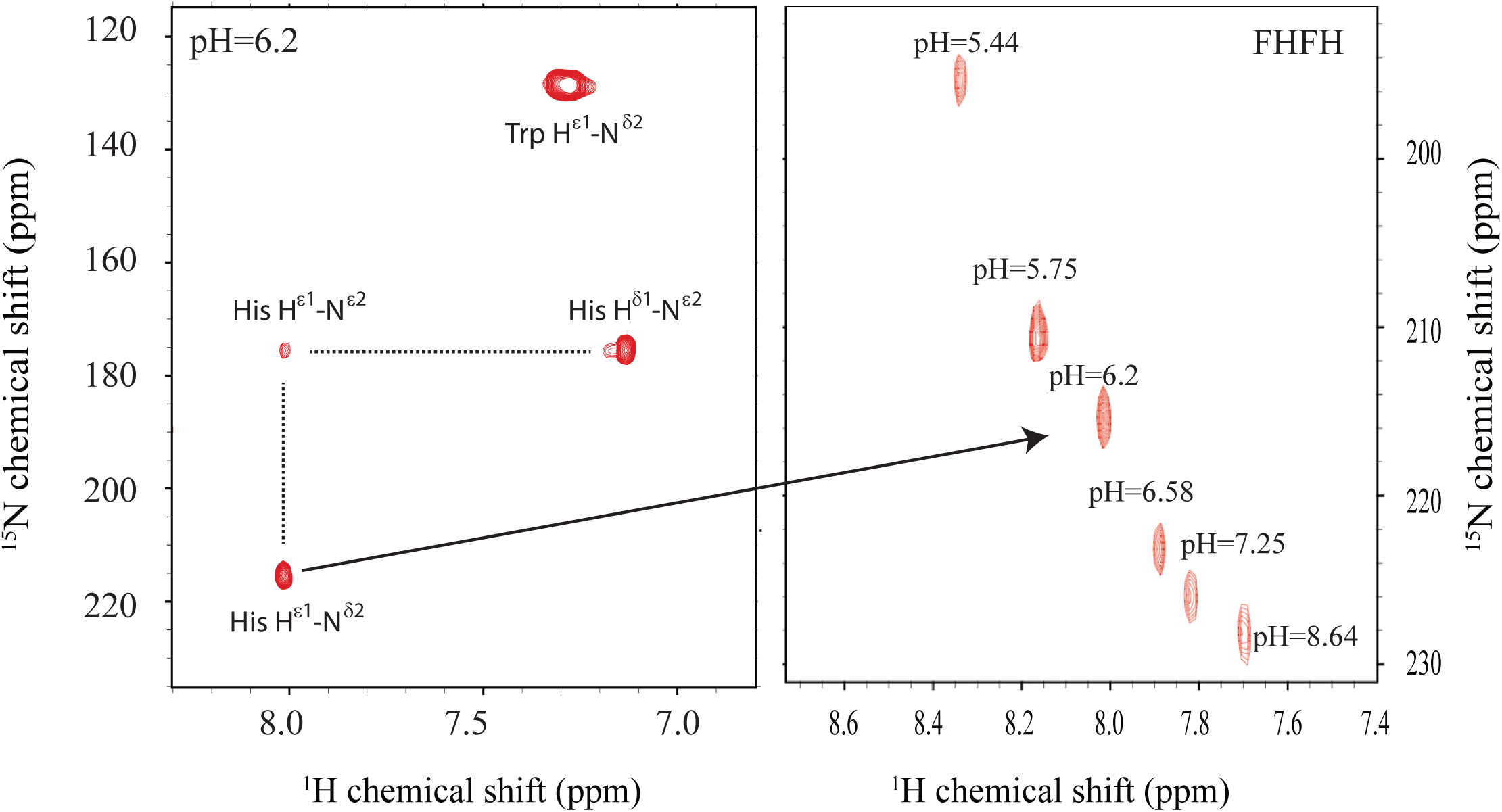
pH Titration of histidine pK_a_s. (A) Multiple bond correlation spectrum of FHFH at pH 6.2 (B) Overlay of multiple-bond correlated H^ε1^-N^δ2^ signals as a function of pH. The ^1^H chemical shifts of this resonance were used to generate the plot in Figure 2A. Fits of the ^15^N shift data give the same result, within error.

## References

1. Kundu, S., Trent, J. T., and Hargrove, M. S. (2003) Plants, humans and hemoglobins. Trends Plant Sci. 8, 387–393

2. Smagghe, B. J., Halder, P., and Hargrove, M. S. (2008) Measurement of distal histidine coordination equilibrium and kinetics in hexacoordinate hemoglobins. in Globins and Other Nitric Oxide-Reactive Proteins, Pt A, Elsevier Academic Press Inc, San Diego. pp 359–378

3. Vallee, B. L., and Williams, R. J. P. (1968) Metalloenzymes - entatic nature of their active sites. Proc. Natl. Acad. Sci. U. S. A. 59, 498–505

4. Zhang, L., Andersen, E. M. E., Khajo, A., Maggliozzo, R. S., and Koder, R. L. (2013) Dynamic factors affecting gaseous ligand binding in an artificial oxygen transport protein. Biochemistry 52, 447–455

5. Zhang, L., Anderson, J. L., Ahmed, I., Norman, J. A., Negron, C., Mutter, A. C., Dutton, P. L., and Koder, R. L. (2011) Manipulating cofactor binding thermodynamics in an artificial oxygen transport protein. Biochemistry 50, 10254–10261

6. Koder, R. L., Anderson, J. L. R., Solomon, L. A., Reddy, K. S., Moser, C. C., and Dutton, P. L. (2009) Design and engineering of an O2 transport protein. Nature 458, 305–309

7. Shikama, K. (1998) The molecular mechanism of autoxidation for myoglobin and hemoglobin: A venerable puzzle. Chem. Rev. 98, 1357–1373

8. Collman, J. P., and Fu, L. (1999) Synthetic models for hemoglobin and myoglobin. Chem. Rev. 32, 455–463

9. Brantley, R. E., Smerdon, S. J., Wilkinson, A. J., Singleton, E. W., and Olson, J. S. (1993) The mechanism of autooxidation of myoglobin. J. Biol. Chem. 268, 6995–7010

10. Anderson, J. L. R., Koder, R. L., Moser, C. C., and Dutton, P. L. (2008) Controlling complexity and water penetration in functional de novo protein design. Biochem. Soc. Trans. 36, 1106–1111

11. Wang, J. H. (1958) Hemoglobin Studies II. A synthetic Material with Hemoglobin-Like Property. J. Am. Chem. Soc. 80, 3168–3169

12. Wang, J. H., Nakahara, A., and Fleischer, E. B. (1958) Hemoglobin Studies I. The Combination Of Carbon Monoxide With Hemoglobin And Related Model Compounds. J. Am. Chem. Soc. 80, 1109–1113

13. Arcon, J. P., Rosi, P., Petruk, A. A., Marti, M. A., and Estrin, D. A. (2015) Molecular Mechanism of Myoglobin Autoxidation: Insights from Computer Simulations. The Journal of Physical Chemistry B 119, 1802–1813

14. Mollan, T. L., Jia, Y., Banerjee, S., Wu, G., Kreulen, R. T., Tsai, A.-L., Olson, J. S., Crumbliss, A. L., and Alayash, A. I. (2014) Redox properties of human hemoglobin in complex with fractionated dimeric and polymeric human haptoglobin. Free Radical Biology and Medicine 69, 265–277

15. Koder, R. L., and Dutton, P. L. (2006) Intelligent design: the de novo engineering of proteins with specified functions. Dalton Trans. 25, 3045–3051

16. Negron, C., Fufezan, C., and Koder, R. L. (2009) Helical Templates for Porphyrin Binding in Designed Proteins. Proteins 74, 400–416

17. Cochran, F. V., Wu, S. P., Wang, W., Nanda, V., Saven, J. G., Therien, M. J., and DeGrado, W. F. (2005) Computational de novo design and characterization of a four-helix bundle protein that selectively binds a nonbiological cofactor. J. Am. Chem. Soc. 127, 1346–1347

18. McAllister, K. A., Zou, H. L., Cochran, F. V., Bender, G. M., Senes, A., Fry, H. C., Nanda, V., Keenan, P. A., Lear, J. D., Saven, J. G., Therien, M. J., Blasie, J. K., and DeGrado, W. F. (2008) Using alpha-helical coiled-coils to design nanostructured metalloporphyrin arrays. J. Am. Chem. Soc. 130, 11921–11927

19. Solomon, L. A., Witten, J., Kodali, G., Moser, C. C., and Dutton, P. L. (2022) Tailorable Tetrahelical Bundles as a Toolkit for Redox Studies. J. Phys. Chem. B 126, 8177–8187

20. Hutchins, G. H., Noble, C. E. M., Bunzel, H. A., Williams, C., Dubiel, P., Yadav, S. K. N., Molinaro, P. M., Barringer, R., Blackburn, H., Hardy, B. J., Parnell, A. E., Landau, C., Race, P. R., Oliver, T. A. A., Koder, R. L., Crump, M. P., Schaffitzel, C., Oliveira, A. S. F., Mulholland, A. J., and Anderson, J. L. R. (2023) An expandable, modular de novo protein platform for precision redox engineering. Proc Natl Acad Sci USA 120, e2306046120

21. Farid, T. A., Kodali, G., Solomon, L. A., Lichtenstein, B. R., Sheehan, M. M., Fry, B. A., Bialas, C., Ennist, N. M., Siedlecki, J. A., Zhao, Z. Y., Stetz, M. A., Valentine, K. G., Anderson, J. L. R., Wand, A. J., Discher, B. M., Moser, C. C., and Dutton, P. L. (2014) Elementary tetrahelical protein design for diverse oxidoreductase functions (vol 9,pg 826, 2013). Nature Chemical Biology 10, 164–164

22. Englander, S. W., Calhoun, D. B., and Englander, J. J. (1987) Biochemistry without oxygen. Anal. Biochem. 161, 300–306

23. Koder, R. L., Valentine, K. G., Cerda, J. F., Noy, D., Smith, K. M., Wand, A. J., and Dutton, P. L. (2006) Native-like structure in designed four helix bundles driven by buried polar interactions. J. Am. Chem. Soc. 128, 14450–14451

24. Berry, E. A., and Trumpower, B. L. (1987) Simultaneous determination of hemes-A, hemes-B, and hemes-C from pyridine hemochrome spectra. Anal. Biochem. 161, 1–15

25. Andrade, M. A., Chacon, P., Merelo, J. J., and Moran, F. (1993) Evaluation of secondary structure of proteins from UV circular-dichroism spectra using an unsupervised learning neural network. Protein Engineering 6, 383–390

26. Huang, S. S., Koder, R. L., Lewis, M., Wand, A. J., and Dutton, P. L. (2004) The HP-1 maquette: From an apoprotein structure to a structured hemoprotein designed to promote redox-coupled proton exchange. Proc. Natl. Acad. Sci. U. S. A. 101, 5536–5541

27. Moffet, D. A., Case, M. A., House, J. C., Vogel, K., Williams, R. D., Spiro, T. G., McLendon, G. L., and Hecht, M. H. (2001) Carbon monoxide binding by de novo heme proteins derived from designed combinatorial libraries. J. Am. Chem. Soc. 123, 2109–2115

28. Trent, J. T., Hvitved, A. N., and Hargrove, M. S. (2001) A model for ligand binding to hexacoordinate hemoglobins. Biochemistry 40, 6155–6163

29. Gardner, A. M., Martin, L. A., Gardner, P. R., Dou, Y., and Olson, J. S. (2000) Steady-state and transient kinetics of Escherichia coli nitric-oxide dioxygenase (flavohemoglobin) - The B10 tyrosine hydroxyl is essential for dioxygen binding and catalysis. J. Biol. Chem. 275, 12581–12589

30. Delaglio, F., Grzesiek, S., Vuister, G., Zhu, G., Pfeifer, J., and Bax, A. (1995) NMRPipe: A Multidimensional Spectral Processing System Based on UNIX Pipes. J. Biomol. NMR 6, 277–293

31. Goddard, T. D., and Kneller, D. G. (2007) Sparky. The University of California, San Francisco

32. Kay, L. E., Keifer, P., and Saarinen, T. (1992) Pure Absorption Gradient Enhanced Heteronuclear Single Quantum Correlation Spectroscopy with Improved Sensitivity. J. Am. Chem. Soc. 114, 10663–10665

33. Pelton, J. G., Torchia, D. A., Meadow, N. D., and Roseman, S. (1993) Tautomeric states of the active-site histidines of phosphorylated and unphosphorylated III(GLC), a signal-transducing protein from escherichia coli, using 2-dimensional heteronuclear NMR techniques. Protein Science 2, 543–558

34. Englander, S. W. (2000) Protein folding intermediates and pathways studied by hydrogen exchange. Annu. Rev. Biophys. Biomolec. Struct. 29, 213–238

35. Brown, M. C., Mutter, A. C., Koder, R. L., JiJi, R. D., and Cooley, J. W. (2013) Observation of persistent -helical content and discrete types of backbone disorder during a molten globule to ordered peptide transition via deep-UV resonance Raman spectroscopy. Journal of Raman Spectroscopy 44, 957–962

36. Balakrishnan, G., Hu, Y., Nielsen, S.B., Spiro, T.G. (2005) Tunable kHz Deep Ultraviolet (193–210 nm) Laser for Raman Applications. Applied Spectroscopy 59, 776–781

37. Ferraro, J. R., Nakamoto, K., Brown, C.V. (1994) Introductory Raman Spectroscopy, 2 ed., Academic Press, San Diego, CA

38. Simpson, J. V., Oshokoya, O., Wagner, N., Liu, J., JiJi, R.D. (2011) Pre-processing of ultraviolet Raman spectra. Analyst 136, 1239–1247

39. Simpson, J. V., Balakrishnan, G., JiJi, R.D. (2009) MCR-ALS analysis of two-way UV resonance Raman spectra to resolve discrete protein secondary structural motifs. Analyst 134, 138–147

40. Mutter, A. C., Norman, J. A., Tiedemann, M. T., Singh, S., Sha, S., Morsi, S., Ahmed, I., Stillman, M. J., and Koder, R. L. (2014) Rational Design of a Zinc Phthalocyanine Binding Protein. Journal of Structural Biology 185, 178–185

41. Ogihara, N. L., Ghirlanda, G., Bryson, J. W., Gingery, M., DeGrado, W. F., and Eisenberg, D. (2001) Design of three-dimensional domain-swapped dimers and fibrous oligomers. Proc. Natl. Acad. Sci. U. S. A. 98, 1404–1409

42. Kim, J., Mao, J., and Gunner, M. R. (2005) Are acidic and basic groups in buried proteins predicted to be ionized? J. Mol. Biol. 348, 1283–1298

43. Englander, S. W., Sosnick, T. R., Englander, J. J., and Mayne, L. (1996) Mechanisms and uses of hydrogen exchange. Curr. Opin. Struct. Biol. 6, 18–23

44. Wang, Y., Purrello, R., Jordan, T., and Spiro, T. G. (1991) UVRR spectroscopy of the peptide bond. 1. Amide-S, A nonhelical structure marker, is a C-alpha-H bending mode. J. Am. Chem. Soc. 113, 6359–6368

45. Halsey, C. M., Oshokoya, O. O., Jiji, R. D., and Cooley, J. W. (2011) Deep-UV Resonance Raman Analysis of the Rhodobacter capsulatus Cytochrome bc(1) Complex Reveals a Potential Marker for the Transmembrane Peptide Backbone. Biochemistry 50, 6531–6538

46. Halsey, C. M., Xiong, J., Oshokoya, O. O., Johnson, J. A., Shinde, S., Beatty, J. T., Ghirlanda, G., Jiji, R. D., and Cooley, J. W. (2011) Simultaneous Observation of Peptide Backbone Lipid Solvation and alpha-Helical Structure by Deep-UV Resonance Raman Spectroscopy. Chembiochem 12, 2125–2128

47. Brown, M. C., Yakuba, R. A., Taylor, J., Halsey, C. M., Xiong, J., JiJi, R. D., and Cooley, J. W. (2014) Bilayer surface association of the pHLIP peptide promotes extensive backbone desolvation and helically-constrained structures. Biophys. Chem. 187-188, 1–6

48. Reedy, C. J., Kennedy, M. L., and Gibney, B. R. (2003) Thermodynamic characterization of ferric and ferrous haem binding to a designed four-alpha-helix protein. Chem. Commun., 570–571

49. Hargrove, M. S. (2000) A flash photolysis method to characterize hexacoordinate hemoglobin kinetics. Biophys. J. 79, 2733–2738

50. Halder, P., Trent, J. T., and Hargrove, M. S. (2007) Influence of the protein matrix on intramolecular histidine ligation in ferric and ferrous hexacoordinate hemoglobins. Proteins 66, 172–182

51. Benson, D. R., Hart, B. R., Zhu, X., and Doughty, M. B. (1995) Design, synthesis, and circular-dichroism investigation of a peptide-sandwiched mesoheme. J. Am. Chem. Soc. 117, 8502–8510

52. Lee, K. H., Kennedy, M. L., Buchalova, M., and Benson, D. R. (2000) Thermodynamics of carbon monoxide binding by helical hemoprotein models: the effect of a competing intramolecular ligand. Tetrahedron 56, 9725–9731

53. Choma, C. T., Lear, J. D., Nelson, M. J., Dutton, P. L., Robertson, D. E., and Degrado, W. F. (1994) Design of a Heme-Binding 4-Helix Bundle. J. Am. Chem. Soc. 116, 856–865

54. Robertson, D. E., Farid, R. S., Moser, C. C., Urbauer, J. L., Mulholland, S. E., Pidikiti, R., Lear, J. D., Wand, A. J., Degrado, W. F., and Dutton, P. L. (1994) Design and Synthesis of Multi-Heme Proteins. Nature 368, 425–431

55. Liu, D. H., Lee, K. H., and Benson, D. R. (1999) Taming the coil: stabilizing a model hemoprotein fold via macrocyclization and peptide helix capping. Chem. Commun., 1205–1206

56. Ghirlanda, G., Osyczka, A., Liu, W. X., Antolovich, M., Smith, K. M., Dutton, P. L., Wand, A. J., and DeGrado, W. F. (2004) De novo design of a D-2-symmetrical protein that reproduces the diheme four-helix bundle in cytochrome bc(1). J. Am. Chem. Soc. 126, 8141–8147

57. Aslund, F., Berndt, K. D., and Holmgren, A. (1997) Redox potentials of glutaredoxins and other thiol-disulfide oxidoreductases of the thioredoxin superfamily determined by direct protein-protein redox equilibria. J. Biol. Chem. 272, 30780–30786

